# Lysosomal swelling and lysis mediate delivery of C3 toxins to their cytoplasmic targets

**DOI:** 10.1101/2022.10.11.511847

**Authors:** Madison Turner, Jonathan Plumb, A. Rod Merrill, Sergio Grinstein, Johannes Westman

## Abstract

Unlike other cholera-like toxins that contain separate binding/translocation and catalytic subunits, C3-like mono-ADP-ribosyltransferases consist of a single subunit that serves both functions. The manner whereby C3 toxins reach the host cell cytoplasm is poorly understood and was addressed in this study by monitoring the fate of fluorescently-labelled C3larvinA. Following binding to the macrophage membrane in a discontinuous, punctate pattern, the toxin was internalized, traversing the endocytic pathway to reach lysosomes. Strikingly, the lysosomes of C3larvinA-treated cells underwent massive swelling over the course of 1-4 hrs. Lysosomal swelling preceded the extensive rearrangement of the cellular F-actin caused by ADP-ribosylation of cytosolic Rho-GTPases. This suggested that lysosome swelling might be required for escape of the toxin into the cytoplasm where the GTPases reside. Accordingly, preventing swelling by osmotic manipulation or by arresting macropinocytosis precluded the F-actin rearrangement. Toxin-induced swelling was associated with leakage of sulforhodamine B and dextran from the lysosomes, implying membrane rupture or activation of mechano-sensitive pores, enabling the toxin itself to reach the cytosol. Finally, comparison of the cellular traffic and actin remodelling activities of C3larvinA with that of two related toxins, C3larvin_trunc_ and Plx2A, highlighted the importance of the N-terminal α_1_-helix for lysosomal swelling and successful infection.

## Introduction

Pathogenic bacteria facilitate the invasion of host tissues through the secretion of virulence factors (Simon *et al*., 2014). These include enzymes that co-opt key pathways of the host cells (Holbourn *et al*., 2006; Yates *et al*., 2006; Fieldhouse and Merrill, 2008). Prototypical examples are the mono-ADP-ribosyltransferase toxins (mARTTs), a diverse family of exoenzymes that facilitate the transfer of an ADP-ribose moiety from a donor NAD^+^ molecule onto a target macromolecule (Holbourn *et al*., 2006; Fieldhouse and Merrill, 2008; Simon *et al*., 2014; Cohen *et al*., 2018). Substrates for this class of toxins vary, and include proteins critical in maintaining host cell homeostasis, such as Gα_s_, actin monomers and, notably, Rho-GTPases (Tainer *et al*., 1999; Han et al., 2001; Simon *et al*., 2014). Despite their shared catalytic function, these toxins employ a range of diverse strategies to target their respective substrates (Holbourn *et al*., 2006; Fieldhouse and Merrill, 2008; Simon *et al*., 2014; Tsuge *et al*., 2015). The structural and functional features underlying this diversity have prompted the division of the mARTT family into subclasses based on their domain organization and catalytic activity (Holbourn *et al*., 2006; Fieldhouse and Merrill, 2008; Simon *et al*., 2014).

Cholera-like toxins (CTs) are a category of mARTTs characterized by shared structural homology to the enzymatic domain of cholera toxin (Fieldhouse and Merrill, 2008; Simon *et al*., 2014). Differences in their catalytic motifs, topology and target substrates have led to further subdivision of this class into the secreted CT, C2 and C3-like subgroups (Fieldhouse and Merrill, 2008; Hottiger *et al*., 2010; Simon *et al*., 2014). CT and C2 proteins are comprised of multiple polypeptide chains that associate to function as heteromeric AB toxins (Tainer *et al*., 1999; Simon *et al*., 2014). The B-domain serves as a receptor-binding protein that facilitates the endocytosis of the enzymatically active A-domain (Fieldhouse and Merrill, 2008; Simon *et al*., 2014). C3-like toxins are a unique subgroup of CTs that function as single-domain enzymes capable of infecting target cells without the use of a B-domain. Instead, an individual polypeptide conducts binding, translocating and catalytic functions to target cytosolic Rho-GTPases (Fieldhouse and Merrill, 2008; Simon *et al*., 2014). ADP-ribosylation by C3 toxins inhibits Rho signalling, resulting in the dysregulation of cytoskeletal actin filaments (Genth *et al*., 2003; Wheeler and Ridley, 2004). Compared to other Rho-targeting enzymes, C3 toxins display high specificity for the RhoA, B and C isoforms, which has promoted their routine use as inhibitors in cell biology studies (Wilde *et al*., 2000; Wheeler and Ridley, 2004; Rohrbeck and Just, 2017). Yet, despite their contribution to understanding Rho-GTPases, little is known about the pathways utilized by C3 toxins to reach their target.

Because –unlike their AB counterparts– C3 toxins lack a dedicated translocating subunit, and were presumed to be internalized non-specifically (Fahrer *et al*., 2010; Rohrbeck and Just, 2017). This view has been revised since these toxins can be selectively internalized by macrophages and monocytes (Fahrer *et al*., 2010). Subsequent investigations suggested that C3 toxins bind to a proteinaceous receptor, and that glycosylation and phosphorylation states of the host cell affect C3 association (Rohrbeck and von Elsner, 2014). C3 toxins can be delivered into early endosomes (Fahrer *et al*., 2010; Fellermann *et al*., 2020); however, the complete pathway used for cellular intoxication remains unknown. Importantly, the mechanism whereby these toxins escape the endocytic pathway to enter the cytosol has yet to be identified.

To investigate these remaining questions, we generated recombinant, fluorescently labelled C3 toxins from *Paenibacillus larvae*, a honeybee pathogen of agricultural importance. Two of the *Paenibacillus* toxins, C3larvinA and Plx2A, have been proven to successfully initiate infection in macrophage cells, while the N-terminally truncated protein C3larvin_trunc_ is not infectious (Krska *et al*., 2015; Ebeling *et al*., 2017; Turner *et al*., 2020). These three proteins, therefore, offer a unique system to investigate the role of the N-terminal helix and the machinery required for cell entry and intoxication.

## Results

### Fluorescently labelled C3larvinA induces the breakdown of cortical actin

C3larvinA was recombinantly expressed in *Escherichia coli* and subsequently conjugated to the amine-reactive fluorophore, DyLight-488. The labelling ratio of the fluorophore to protein was maintained at approximately 1 and size-exclusion chromatography confirmed that the conjugated toxin remained monomeric in solution (data not shown). The treatment of RAW 264.7 macrophages with labelled C3larvinA induced the reorganization of F-actin –visualized by staining with Acti-stain– in a dose-dependent manner (Fig 1A). Submembranous (cortical) F-actin was markedly depleted, while cytoplasmic punctate F-actin structures that were barely detectable in control cells became more apparent. The ratio of these two pools of F-actin, quantified as detailed in Figure 1C, decreased as a function of time after addition of C3larvinA (Fig 1D). The redistribution of F-actin was associated with the appearance of thin protrusions extending from the cell body (Fig 1B). These results are in good agreement with those previously reported for the unlabeled toxin, which induced profound morphological changes in J774A.1 macrophages (Turner *et al*., 2020; Turner *et al*., 2021). By phase contrast, large vacuoles were observed to form in cells treated with C3larvinA (Fig 1B). We concluded that the labelled toxin is biologically active and provides a useful indicator to monitor the internalization pathway.

**Figure 1.**
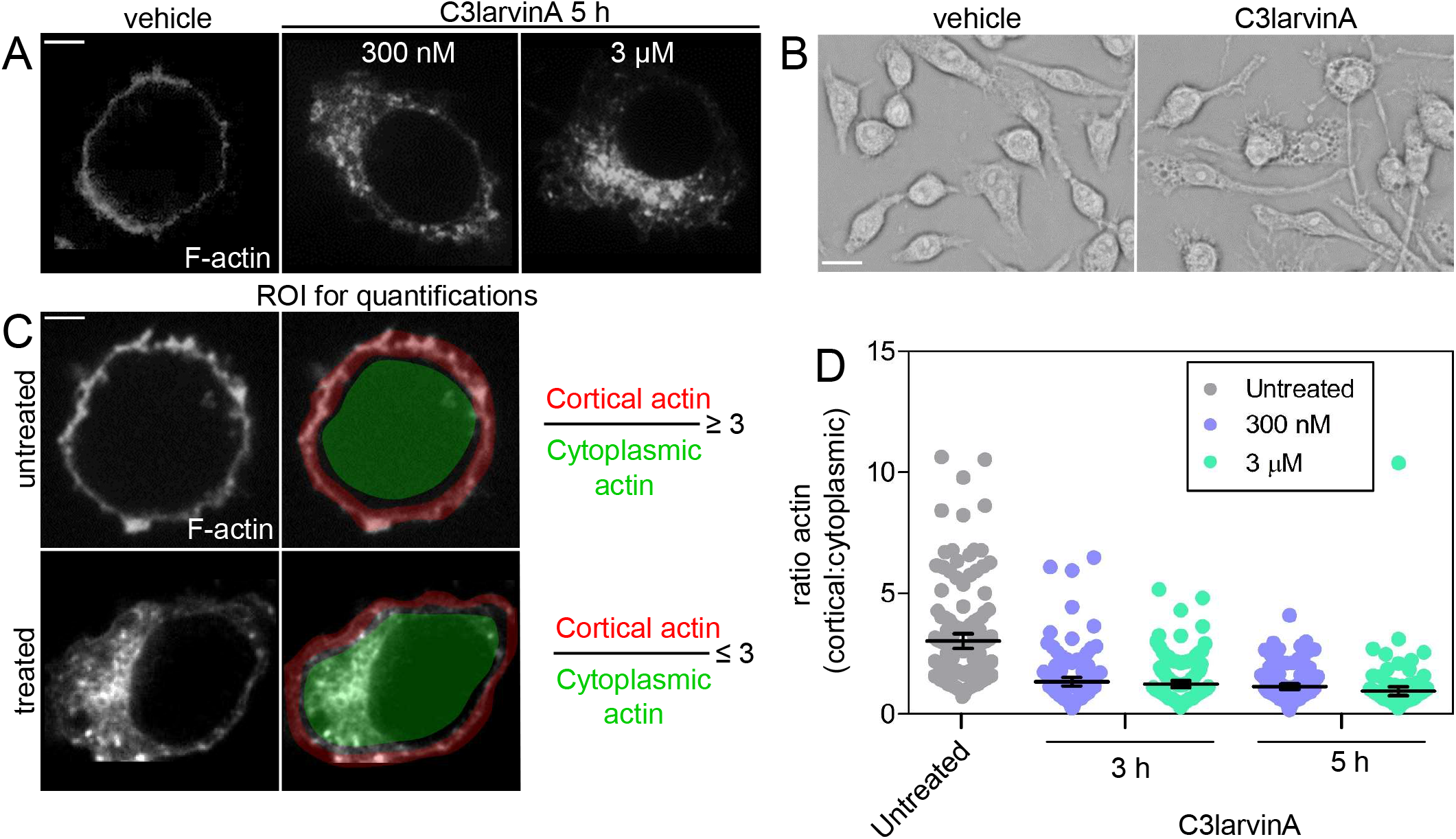
Fluorescently labelled C3larvinA induces the breakdown of cortical actin. (A) Confocal microscopy of RAW 264.7 cells treated without (vehicle only) or with 300 nM or 3 μM C3larvinA-DyLight-488 and incubated for 3 to 5 h at 37°C prior to being fixed and stained with Acti-stain. Scale bar 5 μm. (B) Phase contrast imaging of RAW 264.7 cells treated with 3 μM C3larvinA-DyLight-488 as in (A). Scale bar 25 μm. (C) Description of the procedure used to establish regions or interest (ROIs) for cortical actin (red) and cytoplasmic actin (green) to quantify their ratio. The calculated ratios at 3 h and 5 h of C3larvinA-DyLight-488 treatment are shown in (D). Data points are individual determinations from 3 independent experiments. The mean and SEM are also indicated.

### Cellular traffic of C3larvinA in RAW 264.7 macrophages

To investigate the intracellular fate of C3larvinA, live RAW 264.7 macrophages labelled with various fluorescently tagged organelle markers were monitored using spinning disk confocal microscopy (Fig 2). The cells were initially maintained at 10°C to prevent internalization while being pulsed with C3larvinA for 15 min to promote binding to its putative surface receptors. Following this initial incubation, punctate C3larvinA was visible along the plasma membrane (labelled using concanavalin A), suggesting that a limited number of receptors were engaged or that the receptors had clustered, despite the low temperature (Fig 2A). When the cells were subsequently incubated at 37°C for 5 min, C3larvinA colocalized with transferrin, implying that the toxin was rapidly internalized into early/recycling endosomal vesicles (Fig 2B). After 15 min, the toxin was found to colocalize with the early endosome marker phosphatidylinositol 3-phosphate (PtdIns(3)P), which was detected using PX-RFP, a genetically encoded probe shown earlier to recognize this phosphoinositide specifically (Kanai et al., 2001). Interestingly, at this stage the toxin often appeared in intraluminal vesicles like those known to form in multivesicular bodies. By 30 min, however, the toxin was no longer present in the PtdIns(3)P-positive early compartment (Fig 2C,D). Instead, by this time C3larvinA had reached late endosomes/lysosomes (hereafter referred to collectively as lysosomes), as indicated by its colocalization with lysosome-associated membrane protein 1 (LAMP-1) and with the acidotropic membrane-permeant dye cresyl violet (Ostrowski et al., 2016)(Fig 2E and F). That the toxin-labelled compartment stained with cresyl violet indicated that, at least during the first hours following infection, the lysosomes remain acidic. The acidification may have contributed to the solubilization of the toxin, which at this stage was no longer associated with either the limiting membrane or with intraluminal vesicles, but instead appeared diffuse throughout the lysosomal lumen (Fig 2E and F). It is noteworthy that C3larvinA remained intact for hours inside the lysosomes, despite the proteolytic and acidic nature of this compartment: the molecular weight of the toxin, assessed by immunoblotting, remained unaltered for at least 4 hours (Suppl Fig 1A) and was unaffected when the cells were incubated with a protease inhibitory cocktail or with concanamycin A, a V-ATPase inhibitor known to dissipate lysosomal acidification (Suppl Fig 1B). Therefore, detachment of C3larvinA from the lysosomal limiting membrane cannot be attributed to its proteolysis.

**Figure 2.**
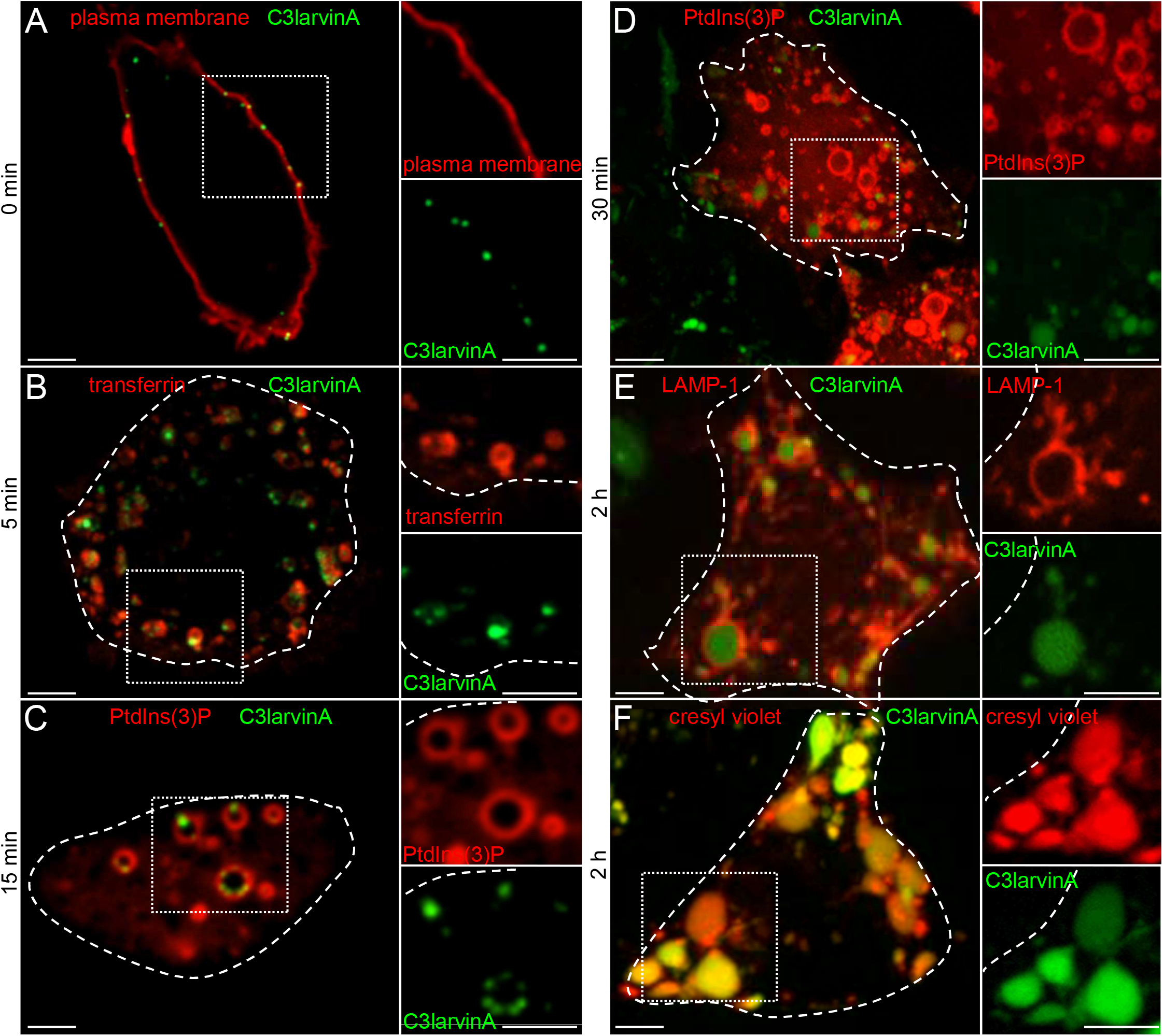
C3larvinA traffic from the plasmalemma to late endosomes/lysosomes. Representative confocal micrographs of live RAW 264.7 cells acquired at the indicated times after exposure to 300 nM of C3larvinA-DyLight-488 (green) at 37°C. (A) Macrophage stained with fluorescently labeled concanavalin A (red). (B) Cells that internalized fluorescently labeled human transferrin (red). (C – E). Cells expressing RFP-tagged PX-RFP, a marker for PtdIns(3)P (C, D) or LAMP-1 (E). (F) Cells were incubated with cresyl violet (red) after 2 h to visualize acidic compartments. All scale bars 5 μm. Images are representative of 3 independent experiments of each type.

Having reached the lysosomes, C3larvinA is unable to return to earlier compartments of the endocytic pathway (Fig 3A) and, unlike some AB toxins, it never reaches the *trans*-Golgi network (Fig 3C) or the endoplasmic reticulum (Fig 3B). Thus, the lysosome appears to be last compartment occupied by the toxin prior to its entry into the cytosol.

**Figure 3.**
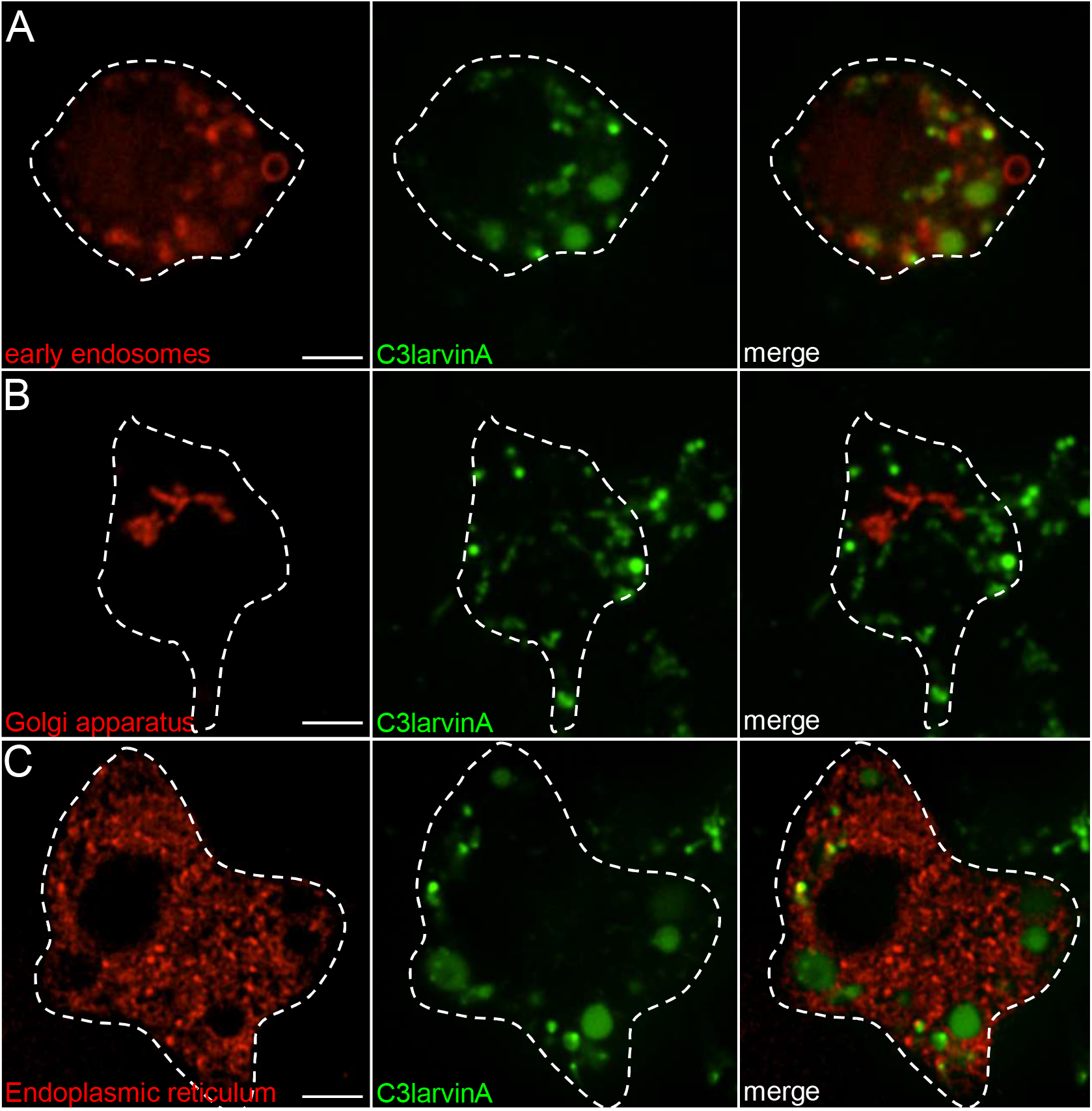
C3larivnA does not co-localize with organelles other than lysosomes at 1 h. Representative confocal micrographs of transfected RAW 264.7 cells expressing RFP- or mCherry-tagged compartment markers for (A) early endosomes, (B) Golgi apparatus, (C) endoplasmic reticulum (all shown in red). Cells were treated with 300 nM C3larvinA-DyLight-488 (green) for 15 min and chased at 37°C for 1 h prior to imaging. All scale bars 5 μm. Images are representative of 3 independent experiments of each type.

### C3larvinA induces swelling of lysosomes

Because the effects of C3larvinA on the cytoskeleton require several hours (e.g., Fig. 1), we monitored its fate beyond the 30-40 min required for it to reach the lysosomes. Extended observation periods revealed that lysosomes containing C3larvinA became progressively swollen; the effect was barely noticeable after 1 h but became more pronounced after 3 h (Fig. 4A). Lysosomal volume, assessed by image reconstruction following acquisition of serial confocal slices, increased >10-fold after 3 h (Fig. 4B). Lysosome swelling was found to be accompanied by a decrease in the total number of toxin-positive lysosomes, suggesting an increase in homotypic fusion events or decreased fission together with lysosome rupture and loss of the labeled toxin, which would similarly decrease the number of detectable lysosomes (Fig 4C).

**Figure 4.**
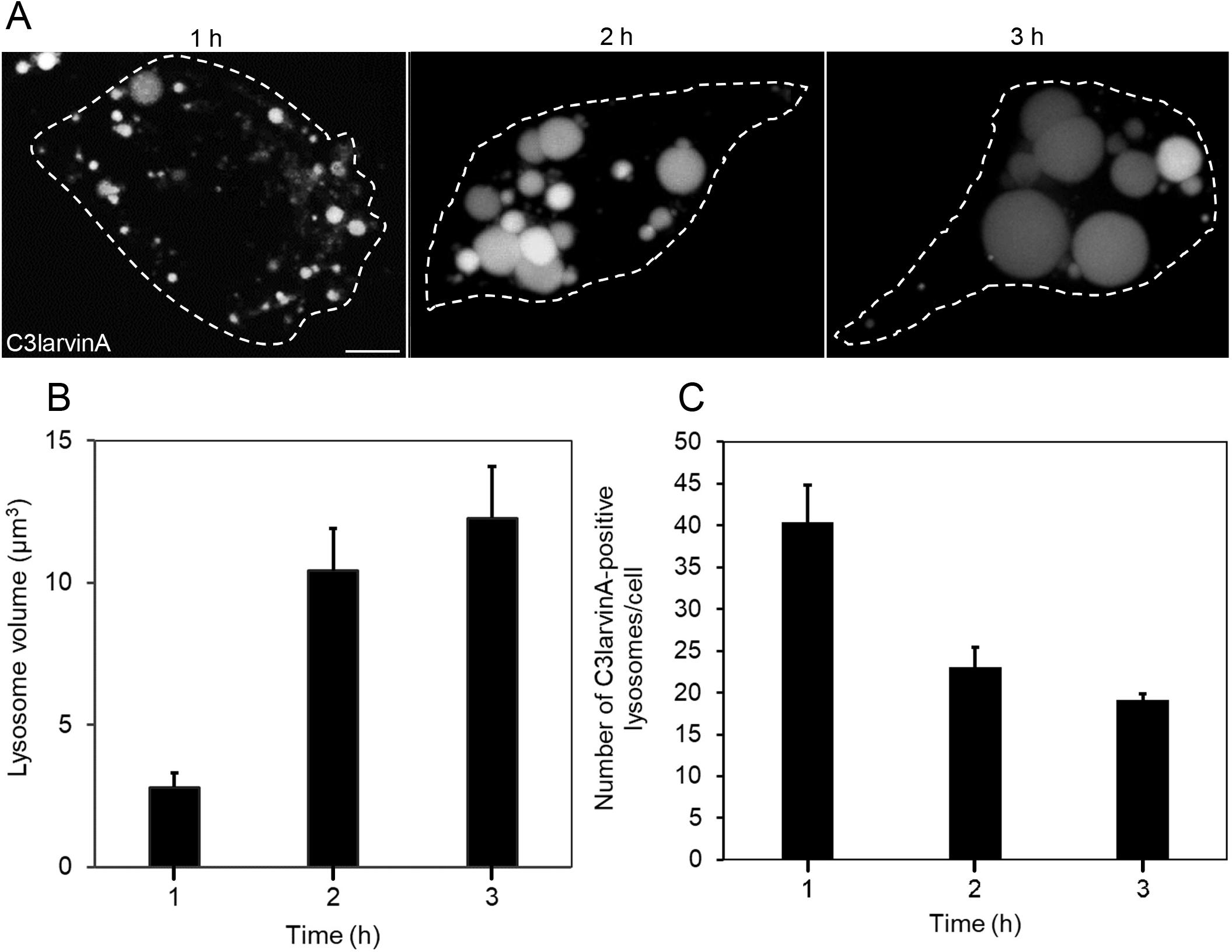
C3larvinA induces swelling of lysosomes. RAW 264.7 cells were treated with 300 nM C3larvinA-DyLight-488 for 15 min and then chased at 37°C for 1 – 3 h. (A) Representative confocal micrographs of live cells showing the distribution of the toxin at 1 – 3 h. Scale bar 5 μm. (B) Lysosomal volume was determined at each time point by assuming the vesicles/vacuoles were spherical (V = 4/3 π r^3^). (C) The number of toxin-positive compartments was determined using confocal sections of the entire height of the cell. Data in B and C are means ± SE of 3 independent experiments, each counting at least 30 cells.

A similar massive swelling and reduced number of lysosomes has been reported in cells treated with inhibitors of PIKfyve, the kinase responsible for conversion of PtdIns(3)P to PtdIns(3,5)P2 (Jefferies et al., 2008; Kim et al., 2014). Indeed, massive vacuoles resembling those seen in cells treated with C3larvinA were also seen when the RAW 264.7 cells were treated with YM201636, a potent and specific PIKfyve inhibitor (Fig. 5A). This raised the possibility that C3larvinA might function as a PIKfyve inhibitor. To evaluate this possibility, we compared the two modes of swelling. In both cases the vacuoles formed were osmotically sensitive since they collapsed when sucrose (200 mM) was added to increase the tonicity of the medium (Fig. 5A). Moreover, as reported in cells treated with PIKfyve inhibitors (Choy et al., 2018), the vacuoles enlarged by C3larvinA did not contain markers of earlier endocytic compartments (Fig. 5B). Despite these similarities, the mechanisms underlying vacuolation in both instances are not identical. As found by others, vacuole enlargement in cells treated with YM201636 was precluded by inhibition of the V-ATPases with concanamycin A (Fig. 5C). By contrast, the C3larvinA-induced swelling was slightly reduced after 2 h and was largely unaffected by concanamycin after 3 h (Fig. 5D). It is therefore unlikely that C3larvinA causes lysosome swelling merely by inhibiting PIKfyve. The partial inhibition of swelling observed in concanamycin-treated cells may explain why contradictory findings have been reported for other C3-like mARTTs in cells treated with bafilomycin A, another V-ATPase inhibitor (Fahrer *et al*., 2010; Rohrbeck *et al*., 2015).

**Figure 5.**
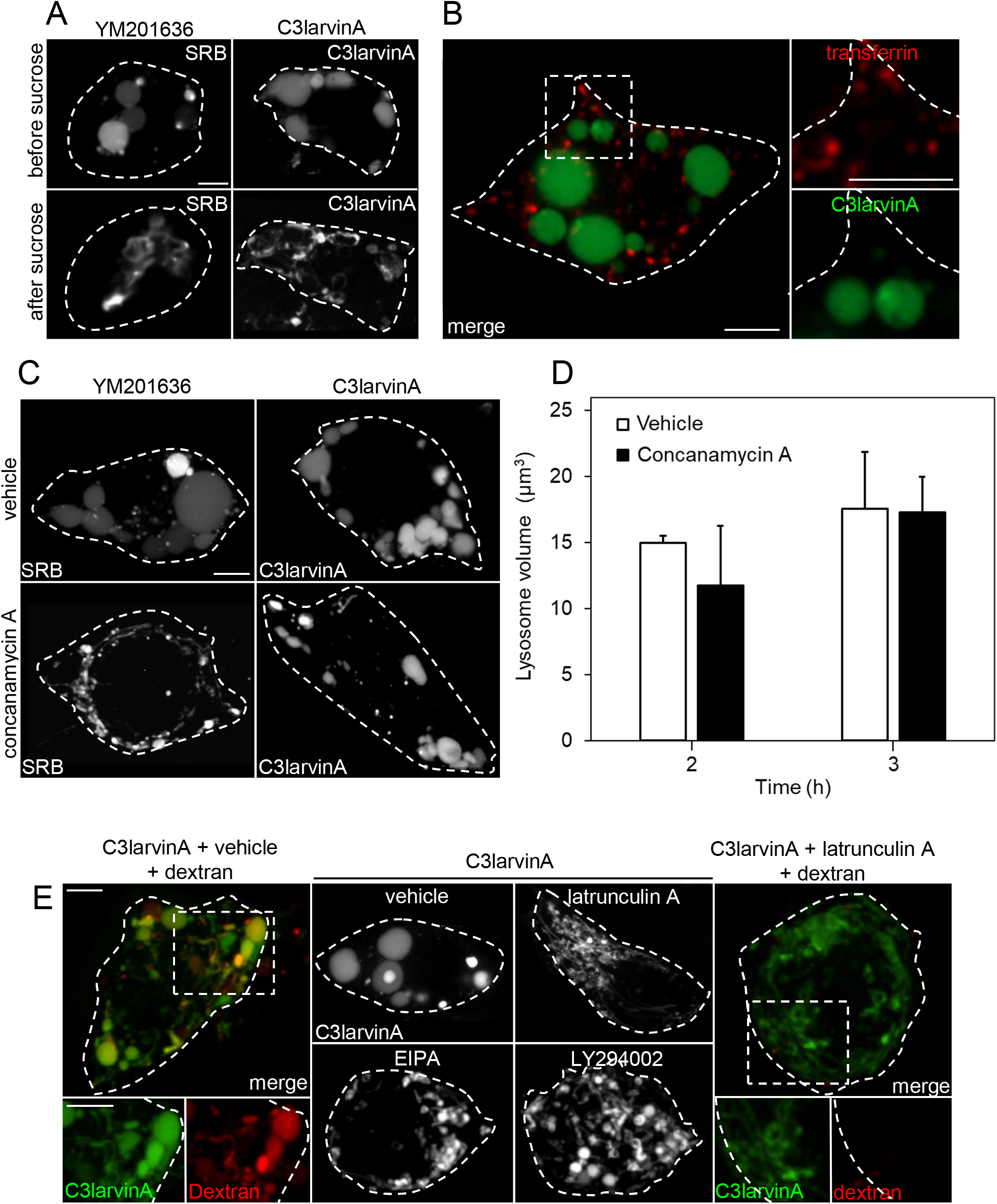
Swelling of lysosomes is sensitive to osmolarity and is dependent on macropinocytosis. (A) Representative micrographs of RAW 264.7 cells treated with YM201636 or C3larvinA-DyLight-488, before and shortly after addition of 200 mM sucrose. (B) Representative micrographs of RAW 264.7 cells treated with C3larvinA-DyLight-488 for 3 h followed by fluorescent transferrin for 5 min. (C) Representative micrographs of RAW 264.7 cells treated with YM201636 or C3larvinA-DyLight-488 followed by concanamycin A. (D) Volume measurements of lysosomes in cells treated with C3larvinA-DyLight-488 with or without concanamycin A for the indicated times. Data are means ± SE of 3 independent experiments, each counting at least 30 cells. (E) RAW 264.7 cells were treated with C3larvinA (green) and 70 kDa dextran (red) in the absence (leftmost panel) or presence of latrunculin A (rightmost panel). Middle panels show the fluorescence of C3larvinA (white) in cells treated with the toxin in the absence (vehicle) or presence of the following inhibitors of macropinocytosis: latrunculin A, EIPA, or LY294002. All scale bars 5 μm. Images in A,B,C and E are representative of 3 independent experiments of each type.

We next studied the contribution of macropinocytosis to the lysosomal enlargement caused by C3larvinA. Macrophages are notorious for their ability to entrap large volumes of extracellular fluid in the form of macropinosomes (Canton, 2018). Inability to process the internalized fluid might explain the effect of C3larvinA on lysosomal volume. The possible role of macropinocytosis was investigated by pulsing RAW 264.7 cells with fluorescent 70 kDa dextran, which is too large to be internalized by conventional modes of endocytosis. First, we confirmed that 70 kDa dextran that entered the cells by macropinocytosis was delivered to the compartment enlarged by C3larvinA (Fig 5E, left panel). To evaluate whether macropinocytosis was required for lysosomal swelling, RAW 264.7 cells were treated with C3larvinA followed by the macropinocytosis antagonists EIPA, latrunculin A, or LY294002, which inhibit Na^+^/H^+^ exchange, actin polymerization and PI3-kinase, respectively. Effective inhibition of macropinocytosis was validated by the suppression of dextran uptake, as illustrated for latrunculin A in Figure 5E, right panel. Importantly, the marked enlargement of lysosomes normally observed in C3larvinA-treated cells was eliminated by EIPA, latrunculin A, or LY294002 (Fig 5, middle panel), implicating macropinocytosis in the swelling process. We concluded that inability to dispose of the fluid and/or membranes internalized by macropinocytosis is responsible, at least in part, for the lysosomal swelling induced by the toxin.

### Lysosomal swelling is critical for the pathogenesis by C3larvinA

The role of lysosome swelling in C3larvinA pathogenesis was investigated next, monitoring the toxin-induced alterations of cortical actin (Fig 6). For these experiments, the toxin was allowed to bind to the cell surface and traffic to the lysosomes, but the subsequent swelling was precluded by increasing the medium osmolarity with hypertonic sucrose, as described for Figure 5A. Preventing lysosomal swelling protected against the effects of the toxin: cortical F-actin remained intact under these conditions and the cytoplasmic F-actin puncta failed to accumulate (Fig 6A and B). Inhibition of lysosome swelling with the macropinocytosis inhibitors EIPA or LY294002 similarly minimized the effects of C3larvinA on actin redistribution (Fig 6C and D). These findings confirm that the continuous delivery of endomembranes and/or their contents is required for C3larvinA-induced pathogenesis in RAW 264.7 macrophages.

**Figure 6.**
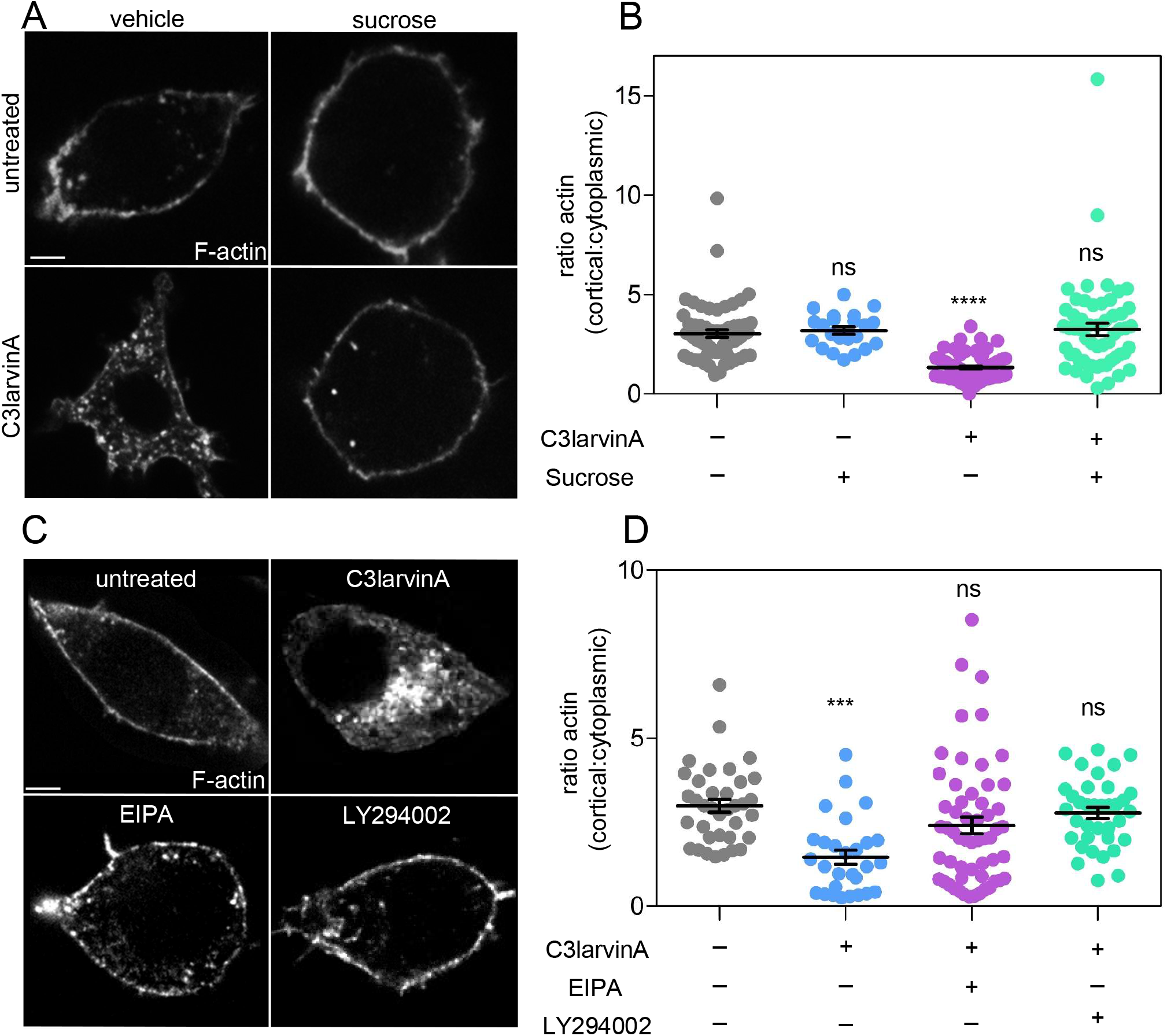
Inhibition of lysosomal swelling prevents the C3larvinA-induced loss of cortical actin. RAW 264.7 cells were treated without or with 300 nM of C3larvinA-DyLight-488 at 37°C for 3 to 5 h in the presence or absence of the indicated inhibitors. Cells were then fixed, and actin was visualized and quantified using Acti-stain-555. (A) Confocal micrographs of cells incubated in isotonic medium (left column) or hypertonic (200 mM) sucrose (right column). Images in A and C are representative of 3 experiments. All scale bars 5 μm. (B) Quantification of the effects of the toxin and of sucrose treatment on F-actin distribution. (C) Representative confocal micrographs of cells treated without (top left) or with toxin in the absence (top right) or presence of EIPA (bottom left) or LY294002 (bottom right). (D) Quantification of the protective effects of EIPA and LY294002 on F-actin redistribution induced by C3larvinA. Data points in B and D are individual cells from 3 independent experiments. The mean and SEM are also indicated.

### C3larvinA-induced swelling causes membrane rupture, enabling the toxin to reach the cytosol

C3larvinA must translocate from the luminal side of lysosomes into the cytosol to ADP-ribosylate its targets (Simon et al., 2014; Turner et al., 2020). Because excessive osmotic swelling can induce defects in the limiting membrane of endocytic organelles (Westman et al., 2018), we investigated whether lysosomal rupture was occurring in C3larvinA-treated cells. The integrity of the lysosomal membrane was assessed by immunostaining the calcium-binding protein ALG-2, which associates with ruptured membranes (Jimenez et al., 2014). Exposure of otherwise untreated (i.e., toxin-free) cells to Gly-Phe-β-naphthylamide (GPN) was used as a positive control. GPN is a membrane-permeant dipeptide and a substrate for cathepsin C. Lysosomal cathepsin C cleaves GPN, generating the membrane-impermeant Phe-β-naphthylamide that accumulates in the lumen, driving osmotically obliged influx of water that causes lysis. As shown in Figure 7A, GPN markedly increased the association of the otherwise diffuse ALG-2 with endomembrane vesicles. Cells treated with C3larvinA also displayed a higher number of ALG-2-stained puncta, suggesting increased endomembrane lysis (Fig 7A). These findings were reproducible and are quantified in Figure 7B.

**Figure 7.**
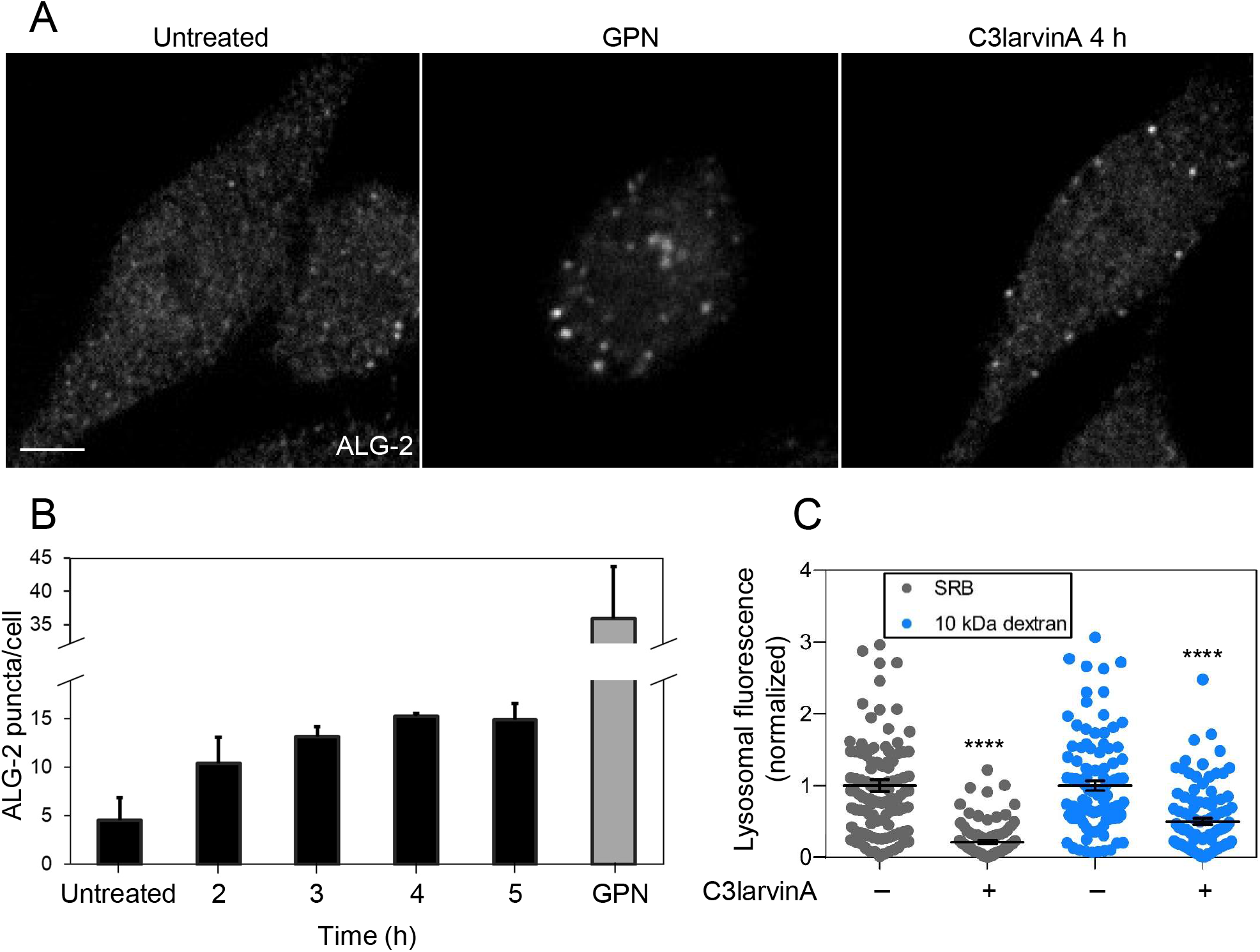
C3larvinA treatment induces rupture of lysosomes. RAW 264.7 cells were either left untreated or were treated with 200 μM of GPN for 60 min or with 300 nM of C3larvinA-DyLight-488 for 20 min and chased for 1 to 5 h at 37°C. Cells were then fixed, permeabilized and stained with anti-ALG-2 antibody. (A) Confocal micrographs, representative of 3 similar experiments. Scale bar 5 μm. (B) Quantification of ALG-2 puncta per cell in the hours following toxin treatment. Puncta were individually counted throughout the entire cell body –reconstructed after serial confocal sectioning– using Fiji. (C) Quantification of lysosome-associated SRB (gray dots) or dextran fluorescence (blue dots) in cells treated with toxin. Cells were pulsed overnight with 25 μg/mL SRB or 50 μg/mL 70 kDa dextran at 37°C. The following day, cells received 300nM C3larvinA-DyLight-488 and were incubated for a further 5 h. Lysosomal fluorescence was quantified in extended focus images reconstructed following serial confocal sectioning of the entire cell.

To directly evaluate membrane lysis, lysosomes were loaded with soluble fluorescent probes that might escape if the membrane ruptures. A pulse-chase protocol was used to load lysosomes selectively with either sulforhodamine B (SRB; MW = 559 g/mol) or with rhodamine dextran, MW 10,000 g/mol, prior to incubation without or with C3larvinA. A marked loss of fluorescence from the lysosomes was observed in cells treated with the toxin for 5 hours, while the soluble probes were retained during the same period in the untreated cells (Fig 7C). Collectively, these results indicate that lysosomal swelling induces permeabilization of the lysosomal membrane, enabling the leakage of molecules as large as 10 kDa. This membrane rupture is the mechanism whereby C3larvinA escapes the endocytic pathway to reach the cytosol.

### Comparing cell entry mechanisms of C3 toxins encoded by Paenibacillus larvae

The N-terminus of C3 toxins has been previously implicated in cell intoxication (Krska *et al*., 2015; Turner *et al*., 2021). The behavior of C3larvinA was therefore compared to two related C3 toxins, C3larvin_trunc_ and Plx2A, which share high sequence homology but vary in N-terminal length (Fig 8A-B) (Ebeling *et al*., 2017; Turner *et al*., 2020). C3larvin_trunc_ and Plx2A were recombinantly expressed in *E. coli* and conjugated to DyLight-488 to a final labelling ratio of approximately 1, as described above for C3larvinA. RAW 264.7 cells were treated with 1 μM of the indicated toxin before being fixed and stained with fluorescent Acti-stain (Fig 8C). Plx2A was found to induce a large redistribution of cortical F-actin, like that caused by C3larvinA, while C3larvin_trunc_-treated cells were indistinguishable from untreated controls (Fig 8D). This agrees with previous literature, which reported large morphological changes in J774A.1 cells treated with Plx2A but not with C3larvin_trunc_ (Krska *et al*., 2015; Ebeling *et al*., 2017).

**Figure 8.**
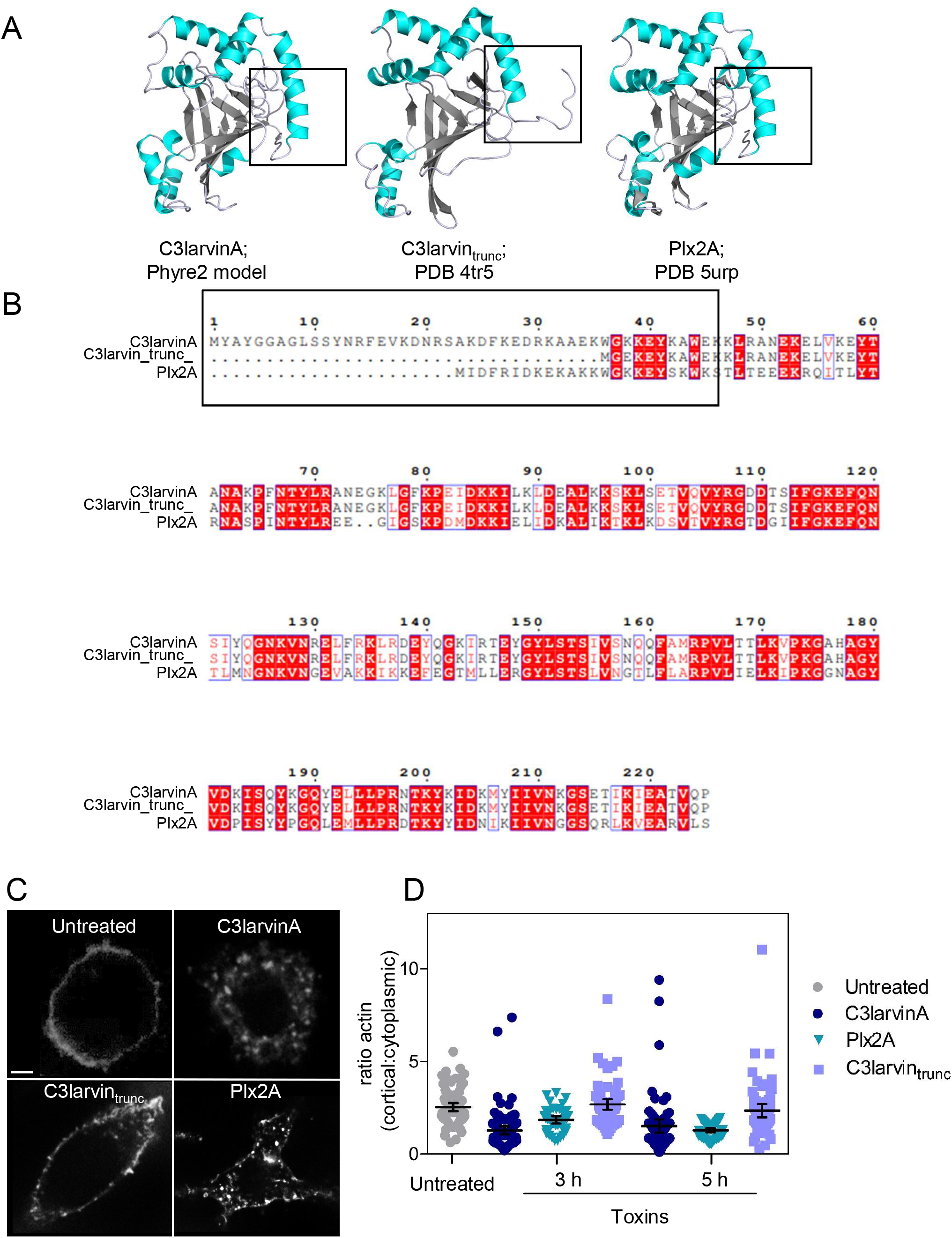
Structural and functional comparison of C3 toxins produced by *Paenibacillus larvae*. (A) Structural comparison of C3larvinA (Phyre 2 homology model), C3larvin_trunc_ (PDB 4tr5) and Plx2A (PDB 5urp). All structures depicted as cartoon, helices coloured cyan, sheets coloured gray and loops coloured light blue; created using PyMOL v1.3. Black boxes highlighting the differences in length of the respective α_1_-helices. (B) Multiple sequence alignment of the three C3-toxins expressed in *P. larvae* created using the Clustal Omega web server and visualized in ESPript. A black box highlights the difference in length of the N-terminal region of the protein sequences. (C) Confocal images of RAW 264.7 macrophage cells treated with 1 μM of the indicated toxin and incubated for 3 to 5 h prior to being fixed and stained with Acti-stain. Scale bar 5 μm. (D) Quantification of the ratio of cortical F-actin to cytoplasmic F-actin from single confocal slices of toxin-treated RAW 264.7 cells after 3 and 5 h. Ratios were determined as in Fig. 1.

Differences in binding, intracellular traffic, or ability to induce lysosomal volume changes could account for the divergent behavior of the toxins. The mechanism underlying their distinctive behavior was investigated next. Like C3larvinA, Plx2A and C3larvin_trunc_ bound to the plasma membrane with a punctate pattern (Fig 9A). Both were internalized via early endosomes (Fig 9B-C) and were delivered to lysosomes (Fig 9D). Interestingly, while Plx2A induced marked lysosomal swelling, resembling the behavior of C3larvinA, C3larvin_trunc_ did not alter the size or shape of the lysosomes. These findings imply that: a) the N-terminus of the toxins is not required for either binding or for internalization but instead contributes to lysosome swelling and b) lysosomal swelling correlates with the biological effects of the toxins, suggesting that the associated rupture is indeed the means whereby the toxins reach their cytoplasmic targets.

**Figure 9.**
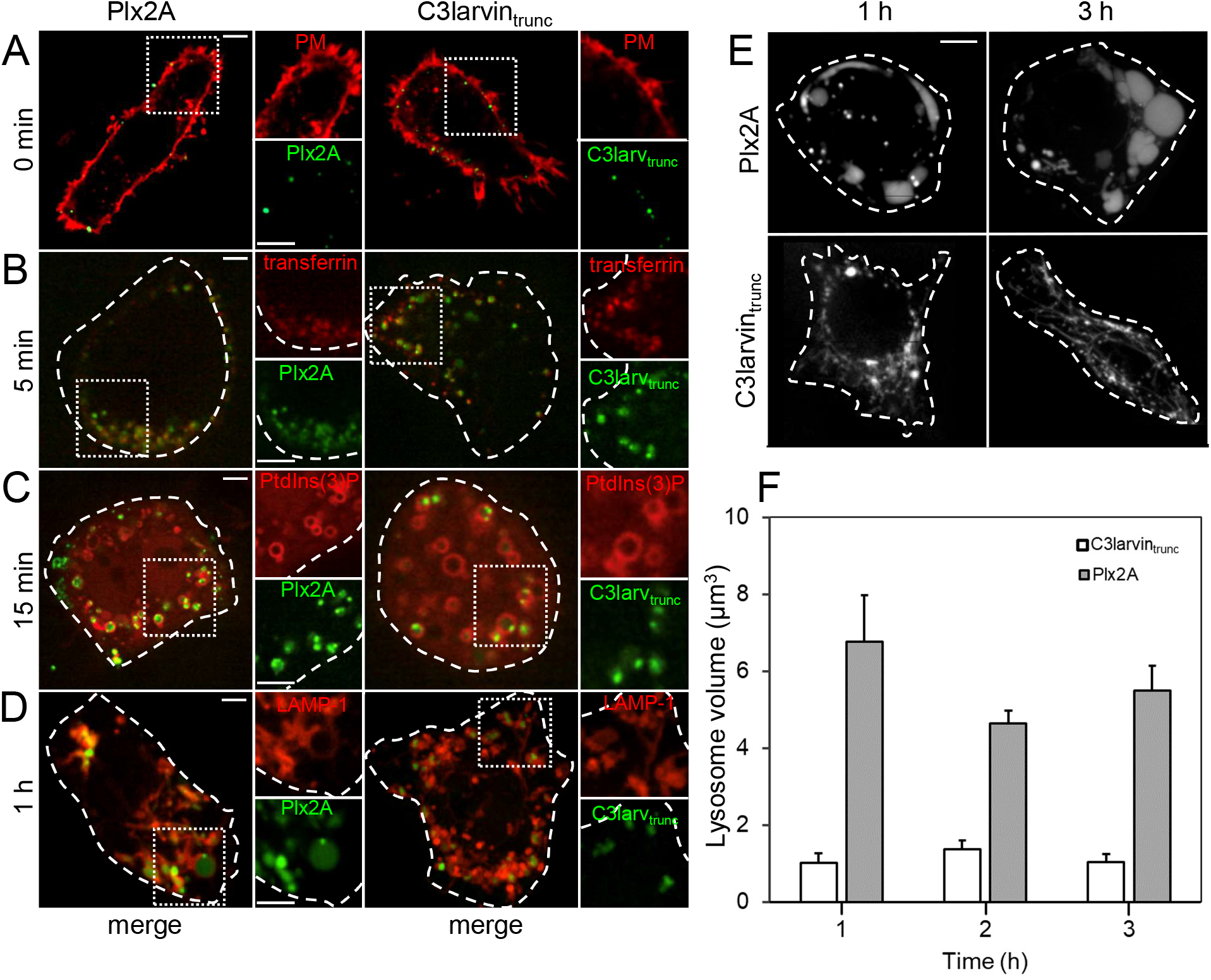
Comparing internalization and toxicity of C3larvin_trunc_ and Plx2A. (A – D) Representative confocal micrographs of live RAW 264.7 cells acquired 0 – 1 h after treatment with either 300 nM Plx2A or C3larvin_trunc_ (green; right and left panels, respectively) at 37°C. (A) Cells stained with fluorescently labeled concanavalin A (red). (B) Cells that internalized the toxins (green) and fluorescently labeled transferrin (red). (C) Cells expressing an RFP-tagged PtdIns(3)P biosensor. (D) Cells expressing RFP-tagged LAMP-1. (E) Confocal micrographs of macrophage cells pulsed with 300 nM of Plx2A (top) or C3larvin_trunc_ (bottom) for 15 min and chased for 1 to 3 h at 37°C to investigate the effect of each toxin on vesicle size and shape. All scale bars 5 μm. (F) Quantification of lysosomal volume at the indicated time points, made as in Fig. 5.

## Discussion

Recombinantly expressed, fluorescently labelled C3larvinA was used as a prototype to investigate the pathway taken by C3-like mARTTs during cell intoxication. The protein was found to bind selectively to the cell membrane before being internalized via early endosomes. We did not attempt to identify the receptor(s) involved, but earlier reports suggested that toxin uptake is reliant upon vimentin (Rohrbeck *et al*., 2014; Rohrbeck *et al*., 2015). We evaluated this hypothesis using bone-marrow-derived macrophages isolated from vimentin knockout mice and found no difference in toxin binding or internalization compared to wild-type macrophages (Suppl Fig. 2); comparable results were obtained for C3larvinA, Plx2A and C3larvin_trunc_. This finding is consistent with the fact that vimentin is a cytoplasmic protein not expected to be exposed exofacially in intact, viable cells. We therefore concluded that vimentin is not essential for intoxication; the main receptor used by C3-like mARTTs remains to be identified.

Like other C3 mARTTs, C3larvinA has been shown to ADP-ribosylate the cytosolic GTPase, RhoA, resulting in the inhibition of Rho signalling (Turner *et al*., 2020). Accordingly, toxin treatment induced the disassembly of cortical F-actin, while increasing intracellular F-actin puncta. The latter may reflect enhanced activity of WASH, a nucleating factor that operates independently of Rho-family GTPases, as reported in cells infected with *Burkholderia* where the glycosyltransferase toxin TcdB similarly targets Rho-GTPases (Walpole et al, 2020). The similarity of the observed phenotypes –breakdown of the cortical actin meshwork with accompanying appearance of cytoplasmic F-actin clusters– suggests that inactivation of Rho-family GTPases, coupled to enhanced WASH activity, underlie actin rearrangement in both instances.

The toxins need to reach the cytosolic compartment to catalyze the ADP-ribosylation of the GTPases. The manner whereby C3 toxins breach the membrane of the endocytic compartments remains poorly understood. In the case of C3larvinA, its internalization was followed by delivery to lysosomes, where it induced their gradual, pronounced swelling. An ensuing membrane rupture and/or opening of mechano-sensitive pores may provide a pathway by which the protein can escape the luminal compartment into the cytosol. Indeed, the amount of SRB and 70 kDa dextran trapped inside lysosomes was found to decrease after toxin treatment. In agreement with this idea, we found that cells could be protected against the toxic effects of C3larvinA when swelling was prevented, whether by increasing medium osmolarity or by inhibiting the ongoing internalization of membranes and entrapped fluid. Macropinocytosis played a particular role in the process, as latrunculin A, LY294002, and EIPA all prevented lysosome swelling when added after the toxin had reached the lysosomes. We speculate that dysregulation of lysosome fission, in the face on continued fusion of upstream vesicular compartments, leads to their enlargement. Homotypic lysosome fusion must also contribute to swelling, since the number of lysosomes decreases in parallel. However, the significant increase in volume:surface ratio that accompanies swelling implies that homotypic fusion is not solely responsible.

The previous characterization of C3 toxins encoded by *P. larvae* uncovered a link between the N-terminal α_1_-helix and successful cell intoxication (Krska *et al*., 2015; Turner *et al*., 2020; Turner *et al*., 2021). This is highlighted by the failure of C3larvin_trunc_ –which has a truncated N-terminal helix– to induce the characteristic morphological changes observed in macrophages treated with the full-length C3larvinA (Krska *et al*., 2015). Cell infection could be restored by extending the N-terminus using residues from another C3 mARTT (Krska *et al*., 2015). Interestingly, C3larvin_trunc_ was nevertheless found to target RhoA *in vitro* and was lethal when expressed in the cytosol of yeast cells (Krska *et al*., 2015). While the truncated version retains catalytic activity, inability to enter the cytosolic compartment is the reason for its lack of toxicity when presented extracellularly. It was unclear, however, how the N-terminal domain effected cell intoxication. In the current study we found that, like C3larvinA and Plx2A –which has an N-terminal helix of intermediate size and effectively alters cytoskeletal structure– C3larvin_trunc_ bound to the macrophage membrane, entered the endocytic pathway, and reached the lysosomes with similar kinetics. Notably, however, C3larvin_trunc_ failed to cause lysosomal swelling, providing further evidence for the crucial role of this event in pathogenesis. It can be concluded that the α_1_-helix contributes to lysosome enlargement, enabling the toxins to enter the cytosol.

Comparative analysis of the N-termini of the three toxins used in this study may provide information regarding their ability to reach the cytosol. Two main points of interest arise from such a comparison: the first is that C3larvinA and Plx2A possess a minimum of 13 additional residues when compared to C3larvin_trunc_. The second is the single difference between the shared sequence of C3larvinA and C3larvin_trunc_ (corresponding to residues 38 and 3, respectively). Where C3larvinA (and Plx2A) possess a Lys residue, C3larvin_trunc_ instead contains an oppositely charged Glu. These differences occur within a 16-residue region that was previously identified to contain multiple conserved motifs postulated to dictate protein structure and function (Lugo and Merrill, 2019). How these residues affect lysosomal swelling and permeabilization is currently unknown, but they are important in directing biological function.

In summary, our findings suggest that osmotic swelling of lysosomes, resulting from fusion of vesicles/vacuoles generated by macropinocytosis and uncompensated by normal physiological fission fosters the leakage of C3 mARTTs to the cytosol, where they reach and covalently modify their targets, notably GTPases. The N-terminal α_1_-helix of the toxins is critical for this effect, although its precise target and mode of action remain to be identified. Additional studies will be required to discern whether the lysosomal compartments are rupturing or if a mechanosensitive pore is responsible for protein leakage. Further analysis is also warranted to elucidate how the N-terminal α_1_-helix initiates the critical lysosome enlargement; identifying its cellular binding partners will undoubtedly provide crucial information.

## Experimental Procedures

### Protein Expression and Purification

C3larvin_trunc_, C3larvinA and Plx2A were all purified as previously described (Krska et al., 2015; Turner et al., 2020; Ebeling et al., 2017). Briefly, protein expression was induced in *E. coli* BL21 λDE3 cells using 1 mM isopropyl β-d-1-thiogalactopyranoside (IPTG) at 37°C for 4 h in the cases of C3larvin_trunc_ and C3larvinA or at 16°C over 18 h in the case of Plx2A. The bacteria were then harvested through centrifugation and resuspended in buffer (0.5 M NaCl, 50 mM Tris-HCl, pH 7.5) containing 120 μM PMSF, 50 μg/mL CHAPS, 100 μg/ml DNase and 1 mM EDTA prior to being lysed in an Emulsiflex C3 high-pressure homogenizer (Avestin Inc., Ottawa, Canada). After a second round of centrifugation, the soluble fractions were incubated with 10 mM MgCl_2_ before isolating the proteins of interest using metal-affinity and size-exclusion chromatography.

Purified protein was exchanged into reaction buffer (0.5 M NaCl, 0.1 M Na_3_PO_4_, pH 7.5) and conjugated to DyLight-488 NHS ester (Thermo Fisher Scientific, Massachusetts, U.S.A.) according to the manufacturer’s instructions, with the following exception: a 1.5 to 2-fold molar excess of dye was used in place of the recommended 8- to 10-fold, to maintain a molar ratio of labelled protein to dye under 1.5. Lastly, the excess dye was removed through dialysis coupled with buffer exchange to isolate conjugated protein in 0.5 M NaCl and 50 mM Tris-HCl, pH 7.5.

### Cell Lines

Experiments were conducted using the RAW 264.7 macrophage cell line (obtained from ATCC, Manassas, VA) derived from a male mouse. The cells were grown at 37°C in an air-CO_2_ (19:1) environment in medium RPMI 1640 (Wisent Inc.) supplemented with 10% (vol/vol) heat-inactivated fetal bovine serum (FBS, Gibco). Prior to experiments, the cells were sparsely plated on glass coverslips inside 12-well tissue culture plates (Corning Inc.) and grown overnight in RPMI 1640 supplemented with 10% FBS.

### Primary Cell Cultures

Bone marrow-derived macrophages were isolated from femurs and tibias of wildtype and vimentin^-/-^ mice (129S-Vim^tm1Cba^/MesDmarkJ; Jackson Laboratories). Briefly, bones from euthanized mice were dissected, immersed in 70% ethanol for 5 min, dried, and the ends removed. The bone marrow was extruded and centrifuged (15,000 x g, 10 sec) into cold phosphate-buffered saline (PBS), washed once in sterile distilled water to lyse red blood cells, and pelleted into cold PBS (500 x g, 10 min). Pellets were suspended in DMEM with 10% FBS, 10 ng/mL M-CSF, 1x antibiotic-antimycotic solution, and plated at a density of 4.0×105 cells per 10 cm-Petri dish for 5-8 days before use.

### Reagents and Plasmids

The following plasmids were described previously: the endoplasmic reticulum marker VAPA-mCherry (Levin-Konigsberg et al., 2019)); the Golgi marker mCh-sialyltransferase was a kind gift from Dr. E. Rodriguez Boulan (Department of Cell Biology, Cornell University); the PtdIns(3)P probe PX-RFP (Kanai et al., 2001); LAMP1-RFP (Martinez et al., 2000; Reddy et al., 2001). Transfection of RAW 264.7 cells was performed using FuGene HD (Promega) according to the manufacturer’s protocols. Antibodies: anti-LAMP-1 hybridoma was from the Developmental Studies Hybridoma Bank; anti-ALG-2 (catalogue #12303-1-AP) was from Proteintech; anti-rabbit secondary antibodies conjugated with Alexa Fluor 488 and Alexa Fluor 647 were from Jackson ImmunoResearch Labs. PBS and Hanks’ balanced salt solution (HBSS) were from Wisent. GPN, ethyl-isopropyl amiloride (EIPA) and cresyl violet were from Sigma-Aldrich; YM201636 and latrunculin A were from Cayman Chemicals; sulforhodamine B was from Molecular Probes; paraformaldehyde was from Electron Microscopy Sciences; Triton X-100 was from Fisher Scientific; concanavalin A conjugated to Alexa Fluor 647 or TMR, 10 kDa dextran and human transferrin-AF647 were from Thermo Fisher. Acti-stain was from Cytoskeleton, Inc. Protease inhibitor cocktail was from Pierce. Concanamycin A was from Abcam and LY294002 from Promega. Sucrose was from BioShop.

### Microscopy

Confocal images were acquired using a spinning disk system (WaveFX; Quorum Technologies Inc.). The instrument consists of a microscope (Axiovert 200M; Zeiss), scanning unit (CSU10; Yokogawa Electric Corporation), electron-multiplied charge-coupled device camera (C9100-13; Hamamatsu Photonics), five-line (405-, 443-, 491-, 561- and 655-nm) laser module (Spectral Applied Research), and filter wheel (MAC5000; Ludl) and is operated by Volocity software version 6.3. Confocal images were acquired using a 63x/1.4-N.A. oil objective (Zeiss) coupled to an additional 1.53 magnifying lens and the appropriate emission filter. Cells imaged live were maintained at 37°C using an environmental chamber (Live Cell Instruments).

Where indicated, epifluorescence images were acquired using an EVOS M5000 Imaging System (Thermo Fisher Scientific). The instrument consists of a high-sensitivity 3.2 MP (2048 × 1536) CMOS monochrome camera with 3.45 μm pixel resolution, three-position chamber (470/525-, 531/593-, 585/624-nm) LED light cubes, and phase contrast imaging mode. Images were acquired using a long working-distance 10×/0.3-N.A air objective (Invitrogen). EVOS M5000 images were analyzed using ImageJ (Fiji 2.1.0/1.53c).

### F-actin Staining

RAW 264.7 cells were treated with 300 nM, 1 μM or 3 μM of toxin for 20 min and chased at 37°C for 3 to 5 hours. For experiments with inhibitors of micropinocytosis, RAW 264.7 cells were treated with 300 nM for 20 min followed by addition of the inhibitor (1 μM latrunculin A, 50 μM LY294002, or 2 μM EIPA, respectively), and chased at 37°C for 3 to 5 hours. Cells were washed twice with PBS before being fixed with 4% PFA for 20 min at room temperature and permeabilized with 0.4% Triton X-100 for 15 min. Samples were incubated for 1 h at room temperature with blocking buffer (2% FBS, 2% BSA) prior to immunostaining for 20 min with a 1:500 dilution of Acti-stain-555 (Cytoskeleton Inc). The ratio of cytoplasmic actin to cortical actin was measured using ImageJ (Fiji 2.1.0/1.53c) from at least 5 random fields per experiment.

### Subcellular Localization of C3larvinA, Plx2A and C3larvin_trunc_

RAW 264.7 cells, wild-type BMDM, or vimentin^-/-^ BMDM were washed with cold PBS and pulsed with 300 nM DyLight-488 labelled toxin for 15 min in RPMI containing 10% FBS at 8°C. To visualize surface-bound toxin the cells were washed in PBS, then incubated for 20 min with 4% cold paraformaldehyde. To monitor endocytosis, cells were pulsed for 20 min with 300 nM DyLight-488 labeled toxin in serum-free RPMI followed by 5 min incubation with 100 μg/mL transferrin AF-488 and washed 2 times with PBS before imaging at 37°C. To visualize other organelles, RAW 264.7 cells were transfected with the indicated organellar markers using FuGene HD 16 hrs prior to treatment with DyLight-488 labeled toxin. For visualization of acidic organelles, 1 μM cresyl violet was added for 5 min, washed two times in PBS and directly imaged by spinning disk confocal microscopy.

### Quantifying Lysosomes and Lysosomal Volume

RAW 264.7 cells were pulsed with 300 nM of C3larvinA-DyLight-488 for 15 min and chased at 37°C. C3larvinA-retaining lysosomes were analyzed acquiring confocal sections encompassing the entire height of entire cells (at least 15 mm) and quantified using the “Analyze Particles” tool in Fiji. To measure lysosomal volume, cells were pulsed with 300 nM of C3larvinA-DyLight-488 for 15 min and chased at 37°C for 1-4 h in the presence of vehicle (DMSO) or the indicated agents (2 μM YM201636; 1 μM concanamycin A; 2 μM EIPA; 50 μM LY294002, or 200 mM sucrose). The diameter of lysosomes was measured at indicated time points using Fiji. The volume of lysosomes was calculated assuming that lysosomes were spherical (volume = 4/3 π r^3^).

### Assessment of lysosomal leakage during swelling

RAW cells were incubated overnight with either 25 μg/mL sulforhodamine B or 50 μg/mL TMR dextran (10 kDa). On the following day, cells were rinsed twice with RPMI-FBS, and 0.5 mL RRPMI-FBS was added per well with or without 300 nM C3larvinA-DyLight-488 and incubated for 5 hours. Serial confocal images were captured at 0.25 μm intervals and used to generate extended focus images to assess the leakage of the fluorophores from the lysosomes.

### ALG-2 Immunostaining

RAW 264.7 cells were grown on glass coverslips as described above and incubated with 300 nM of C3larvinA for 20 min before being washed with PBS and supplemented with fresh medium. Lysis induced by 200 μM GPN was used as a positive control for lysosomal damage. At the indicated time points, samples were washed with PBS and fixed with 4% PFA for 20 min at room temperature. Samples were washed twice in PBS, permeabilized with 0.2% Triton X-100 (15 min, room temperature), and incubated with blocking buffer (2% FBS, 2% BSA) for 60 min at room temperature. Samples were immunostained with rabbit anti-ALG-2 antibody (1:100 in blocking buffer, 60 min) and a fluorescent secondary antibody (1:1000 in blocking buffer, 30 min).

### SDS-PAGE

RAW 264.7 cells were seeded in 12-well plates and grown for two days to 90% confluency. Macrophages were pre-incubated with vehicle or 1X protease inhibitor cocktail for 60 min at 37°C. C3larvinA-DyLight-488 was added to RAW 264.7 cells as described above. Concanamycin A was added after 15 min incubation with C3larvinA-DyLight-488. After 2 hrs the supernatant was withdrawn and 200 μL Laemmli sample buffer (containing 2-mercaptoethanol at 95°C) was added. the cells scraped and denatured for 10 min at 95°C. All samples were separated on 12% sodium dodecyl-sulfate polyacrylamide gels, and the fluorescent C3larvinA-DyLight-488 was visualized using a ChemidDoc XRS+ (Bio-Rad).

### C3larvinA homology model

The C3larvinA homology model was built as previously described (Turner et al., 2020). Briefly, the crystal structure of Plx2A (PDB: 5urp), which was resolved to 1.65 Å and shares 55% sequence identity to C3larvinA, was used as a template to model the C3larvinA structure in Phyre2 (Kelley et al., 2015). The homology model was reported with 100% confidence.

## QUANTIFICATION AND STATISTICAL ANALYSIS

### Image Analysis and Statistics

Image handling, quantification, and analysis of fluorescence images were performed using Volocity 6.3 software (PerkinElmer Inc), Imaris 9.5.1 software (Oxford Instruments), or ImageJ (Fiji v. 2.0.0-rc-65/1.51w) software. The figure legends describe the exact number of independent replicates that were analyzed in each experiment. A total of at least 30 cells were quantified for each condition, and the data presented includes all measured data points for all experiments, including means ± SEM, or bar plots. Statistical analysis was performed using the means of 3-5 individual experiments, and statistical significance was determined using unpaired t-test, non-parametric Mann-Whitney U-test, and one-way analysis of variance (ANOVA) (Tukey’s, Dunnett’s, or Sidak’s test) with Prism 8 (GraphPad Software), with p < 0.05 considered significant.

## Supplemental figure legends

**Suppl. Fig 1.**
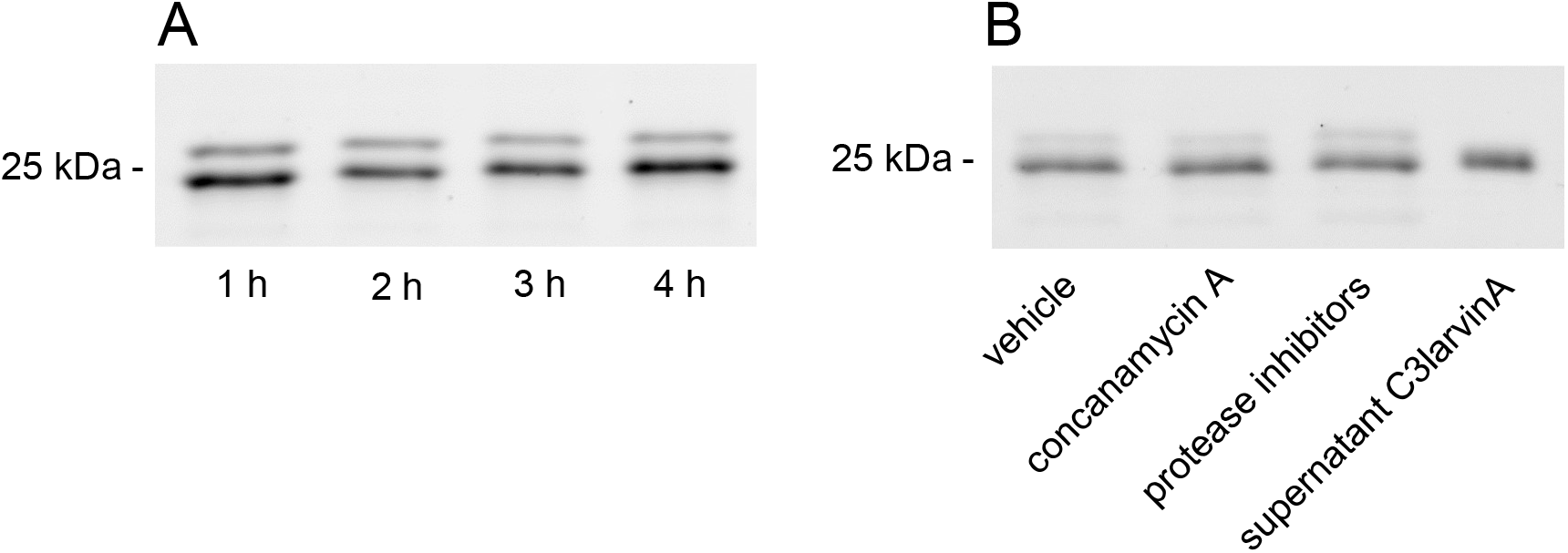
C3larvinA-DyLight-488 remains intact in lysosomes of RAW264.7 macrophages. (A) RAW 264.7 macrophages were treated for the indicated times with C3larvinA-DyLight-488 and then solubilized and analyzed by SDS-PAGE. The fluorescence of the toxin was visualized using a ChemidDoc XRS+ (B) SDS-PAGE of RAW 264.7 macrophages treated for 1 h with C3larvinA-DyLight-488 in the presence of DMSO (vehicle), concanamycin A, or protease inhibitor cocktail. Non-internalized C3larvinA-DyLight-488 collected from the supernatant served as positive control (rightmost lane).

**Suppl. Fig 2.**
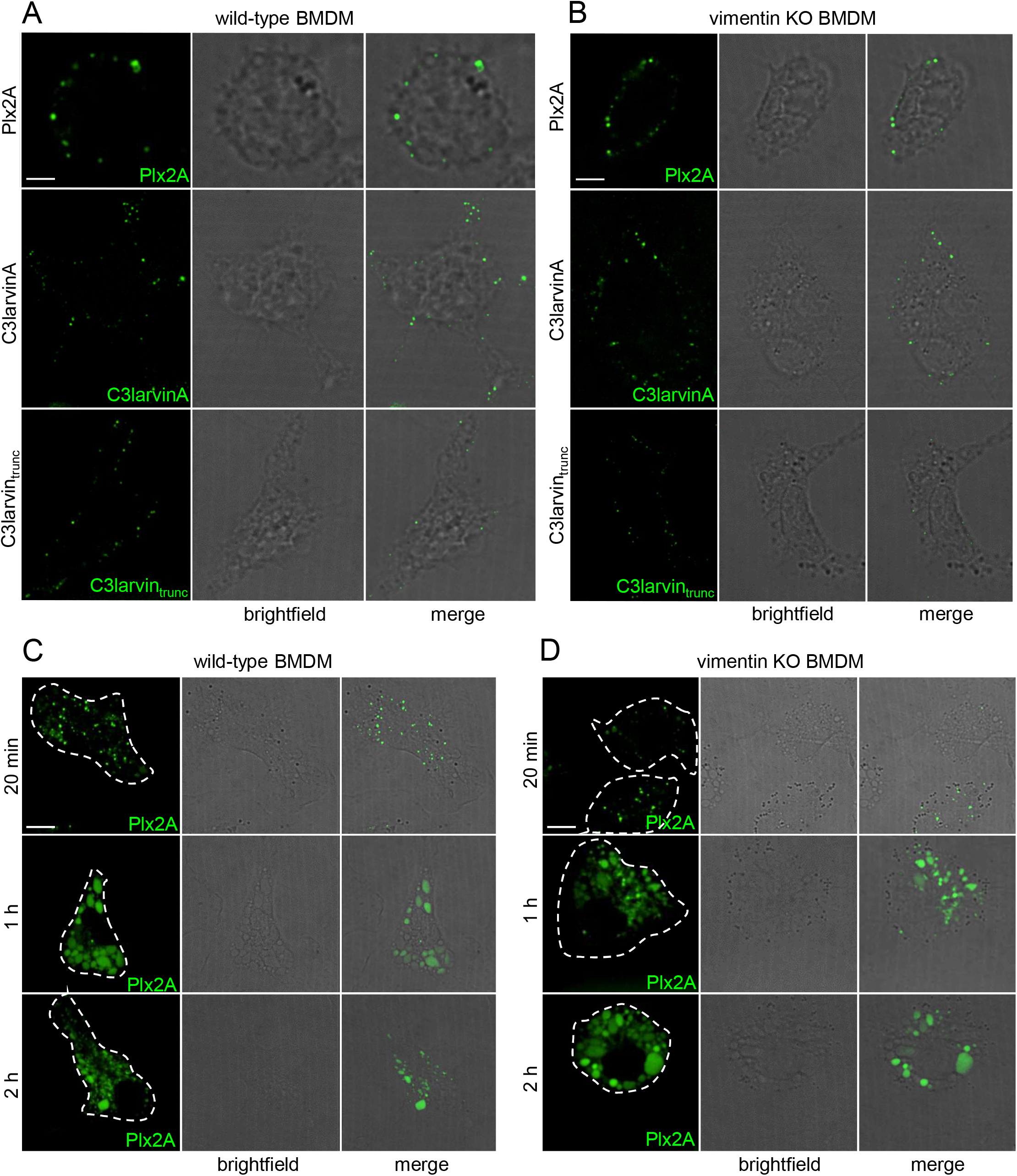
Vimentin is not required for uptake of C3-like toxins or swelling of lysosomes in BMDM macrophages. Macrophages derived from the bone marrow of wild-type or vimentin KO mice were pulsed with C3larvinA-DyLight-488 for 15 min and chased for the indicated time. (A) Representative images of Plx2A, C3larvinA, and C3larvin_trunc_ bound to the surface of wild-type (A) or vimentin KO (B) BMDM. Plx2A induces lysosome swelling in both wild-type (C) and vimentin KO (D) BMDM.

